# Continuous Hypoxia Reduces Retinal Ganglion Cell Degeneration in a Mouse Model of Mitochondrial Optic Neuropathy

**DOI:** 10.1101/2022.09.15.508111

**Authors:** Alexander M. Warwick, Howard M. Bomze, Luyu Wang, Mikael Klingeborn, Ying Hao, Sandra S. Stinnett, Sidney M. Gospe

## Abstract

**Purpose:** To test whether continuous hypoxia is neuroprotective to retinal ganglion cells (RGCs) in a mouse model of mitochondrial optic neuropathy.

**Methods:** RGC degeneration was assessed in genetically modified mice in which the *floxed* gene for the complex I subunit NDUFS4 is deleted from RGCs using *Vlgut2*-driven Cre recombinase. Beginning at postnatal day 25 (P25), *Vglut2-Cre;ndufs4^loxP/loxP^* mice and control littermates were housed under hypoxia (11% oxygen) or were kept under normoxia (21% oxygen). Survival of RGC somas and axons was assessed at P60 and P90 via histological analysis of retinal flat mounts and optic nerve cross sections, respectively. Retinal tissue was also assessed for neuroinflammation using Western blot and confocal microscopy.

**Results:** Consistent with our previous characterization of this model, at least one-third of RGCs had degenerated by P60 in *Vglut2-Cre;ndufs4^loxP/loxP^* mice remaining under normoxia. However, continuous hypoxia resulted in complete rescue of RGC somas and axons at this time point, with normal axonal myelination observed on electron microscopy. Though only partial, hypoxia-mediated rescue of complex I-deficient RGC somas and axons remained significant at P90. Hypoxia prevented reactive gliosis at P60, while the retinal accumulation of Iba1-positive mononuclear inflammatory cells was not substantially reduced.

**Conclusions:** Continuous hypoxia achieved dramatic rescue of early RGC degeneration in mice with severe mitochondrial dysfunction. Although complete rescue was not durable to P90, our observations suggest that investigating the mechanisms underlying hypoxia-mediated neuroprotection of RGCs may identify useful therapeutic strategies for optic neuropathies resulting from less profound mitochondrial impairment, such as Leber Hereditary Optic Neuropathy.

## INTRODUCTION

Mitochondrial dysfunction is an underlying contributor to a variety of optic neuropathies, some of which arise from heritable mutations.^1,2^ The prime example of a mitochondrial optic neuropathy is Leber Hereditary Optic Neuropathy (LHON), which is characterized by rapidly progressive, profound vision loss in both eyes due to degeneration of retinal ganglion cells (RGCs), manifesting most often in adolescents and young adults.^3^ LHON is among the most common mitochondrial diseases, with a reported prevalence of 1:30,000 to 1:50,000.^4-6^ With rare exception,^7,8^ it is caused by mutations in the mitochondrial DNA (mtDNA), resulting in partial loss of function of mitochondrial complex I, a large protein complex localizing to the mitochondrial inner membrane that is responsible for entry of electrons from NADH into the electron transport chain. Complex I is composed of seven mtDNA-encoded subunits and at least 37 nuclear-encoded subunits.^9^ Loss of complex I activity impairs the oxidative ATP-generating capacity of cells,^10^ and perhaps even more importantly results in the leakage of electrons and the formation of deleterious reactive oxygen species.^11^

LHON has been challenging to model in mammals due to the technical difficulty of manipulating the mitochondrial genome. Previously characterized animal models have involved *in vivo* delivery of mutant complex I genes to RGCs by adeno-associated virus or use of genetically modified mouse lines. These models have demonstrated RGC degeneration, but with a latency of at least six months and in some cases one to two years, limiting their utility for rapid screening of potential therapeutic strategies.^12-15^ We recently adapted a mouse model of Leigh syndrome, a severe systemic mitochondrial disease, in order to develop a genetically modified mouse line with much more rapid RGC degeneration than observed in these LHON models.^16^ Deletion of the nuclear gene *ndufs4*, which encodes an accessory complex I subunit mutated in some forms of Leigh syndrome, severely destabilizes complex I and decreases its enzymatic activity by >50% in the retina and other tissues.^17-20^ This severe compromise of complex I function produces a more profound phenotype than seen in LHON, with germline *ndufs4* knockout mice exhibiting a rapidly progressive myoencephalopathy that results in death by around postnatal day 50 (P50).^18^ In order to generate a model of mitochondrial optic neuropathy in which RGC degeneration could be studied over a longer period, we used Cre recombinase driven by the vesicular glutamate transporter V-Glut2 to delete *floxed ndufs4* alleles from RGCs. With complex I dysfunction induced in all RGCs but in only a subset of central nervous system neurons, the *Vglut2-Cre;ndufs4^loxP/loxP^* conditional knockout mice survive nearly twice as long as germline knockouts and manifest a rapid degeneration of RGCs that begins ∼P45 and progresses to loss of more than half of RGCs by P90.^16^ The onset of acute RGC loss just after the mice reach sexual maturity is similar to many human cases of LHON and supports the use of this mouse line as a preclinical model for the disease.

The germline *ndufs4^-/-^*mouse line has been very well-studied in recent years, and several laboratories have reported pharmacological interventions that may prolong the lifespan of these mice.^21-25^ Aside from gene therapy that restores *ndufs4* expression,^26,27^ the most successful therapy described in this mouse model has been to rear the mutant mice under continuous normobaric hypoxia. Mootha and colleagues have reported that housing the mice at an 11% O_2_concentration results in prolongation of the median lifespan to 270 days while also reducing neurological dysfunction.^28,29^ It has been proposed that reduction of the O_2_ tension at the tissue level in this mouse model is critical in relieving the burden of reactive oxygen species formation.^30^

As the ocular phenotype has not been reported in hypoxia-treated germline *ndufs4^-/-^*mice, we were interested to explore the potential therapeutic value of hypoxia in our model of RGC-specific *ndufs4* deletion. Here we report that an 11% O_2_ environment achieved a striking 100% rescue of RGC soma and axon degeneration at P60, along with a reduction of some signs of retinal neuroinflammation. While incomplete, therapeutic efficacy remained robust at P90, indicating that further exploration of the mechanism(s) underlying hypoxia-mediated rescue of complex I-deficient RGCs may identify promising therapeutic targets for patients with mitochondrial optic neuropathies, including LHON.

## METHODS

### Animals

All animal experiments adhered to the ARVO Statement for the Use of Animals in Ophthalmic and Vision Research, following a protocol approved by the Institutional Animal Care and Use Committee of Duke University. *Vglut2-Cre;ndufs4^loxP/loxP^* mice and control littermates were generated as previously described^16^ and maintained on a C57Bl/6J background.

### Continuous Hypoxia

For hypoxia experiments, mouse cages were kept within a hypoxia chamber (A-Chamber animal cage enclosure, BioSpherix, Ltd., Parish, NY) which lowers the ambient PO_2_ by pumping in nitrogen gas to displace the oxygen. Chamber PO_2_ level was set to a constant 11% O_2_. For all experiments, mice born under normoxia were placed into the hypoxia chamber at P25 and maintained there until P60 or P90, under a 12-hour light-dark cycle. Cages with control littermates were maintained on their rack under a normoxic 21% O_2_ concentration. At the indicated time points, the mice were removed from the hypoxia chamber and rapidly euthanized, followed by tissue harvesting for histology.

### Antibodies

The following antibodies were used for immunofluorescence experiments: rabbit polyclonal anti-RBPMS1 (1:500; Novus, NBP2-20112, Lot#130-96), rabbit polyclonal anti-Iba1 (1:1000; Fujifilm Wako Chemicals Corp., 019–19741), and rabbit polyclonal anti-GFAP (D1F4Q) XP (1:200; Cell Signaling Technology, Lot#12389). For Western blot analysis of retinal lysates, the rabbit polyclonal anti-GFAP (D1F4Q) XP antibody was used at a dilution of 1:1000, and mouse monoclonal anti-β-actin (1:1000; Santa Cruz, sc-47778, Lot#D0615) was used as a loading control. Secondary antibodies against the appropriate species conjugated to Alexa Fluor 488 or Alexa Fluor 568 (immunofluorescence experiments, 1:500 dilution) or Alexa Fluor 680 (Western blot experiments, 1:20,000 dilution) were purchased from Invitrogen. Cell nuclei were stained using Hoechst 33342 (1:1000, Thermo Fisher Scientific).

### Histological Techniques

Immunohistochemistry experiments were performed as previously described.^16,31^

Briefly, posterior eyecups obtained from euthanized mice were fixed for 1 h in 4% paraformaldehyde. Retinal flat mounts were then prepared by isolating the retinas, blocking in 5% goat serum in PBS with 0.3% Triton X-100, incubating with anti-RBPMS1 primary antibody in block for 5 days at 4°C, and incubating with anti-rabbit Alexa Fluor 488 in block overnight at 4°C. The retinas were then washed and placed on glass slides with the RGC layer facing up, and four radial cuts were made from the edge to the equator of each retina to achieve flattening prior to mounting.

Retinal cryosections were prepared by cryoprotecting fixed eyecups in 30% sucrose and then embedding them in optimal cutting temperature (OCT) medium (Tissue-Tek, Sakura Finetek). Retinal cross-sections, 20μm in thickness, were collected using a cryostat microtome (Microm HM 550, Thermo Fisher Scientific). Sections were rehydrated with PBS, blocked in 5% goat serum in PBS with 0.3% Triton X-100, and then incubated in primary antibody in the same blocking solution overnight at 4°C. Sections were washed and incubated with appropriate secondary antibody conjugated to Alexa Fluor 488 overnight at 4°C, and washed prior to mounting.

All samples were mounted with Immu-Mount (Thermo Fisher Scientific) under glass coverslips. Images were acquired using a Nikon Eclipse Ti2 inverted confocal microscope, a CFI Plan Fluor 60× (oil) objective, and an A1 confocal scanner controlled by NIS-Elements software (Nikon). For RGC soma quantification in retinal flat mounts, images of 45,000 μm^2^ in area were obtained in each quadrant at locations of 0.5, 1.0, and 1.5 mm from the optic nerve head, and RGC somas were manually counted using the Cell Counter plugin for Fiji.^32^ RGC soma density was averaged among the four quadrants at each distance from the optic nerve head. For quantification of Iba1-positive cells, retinal cross sections were imaged along their entire length, and the number of labeled cells within the ganglion cell and inner plexiform layers was quantified in 3–4 retinal sections per sample (taken through the optic nerve head), then averaged. For quantification of GFAP-positive radial processes of Müller glia, 45,000 μm^2^ images were acquired at a 500-μm distance to either side of the optic nerve head for 3 sections per sample.

The number of positive radial processes present at the inner nuclear layer/inner plexiform layer junction was counted for each section and averaged. To assess RGC axons, mouse optic nerves were obtained from euthanized mice that had undergone transcardial perfusion with 4% PFA and then post-fixed for an additional two hours in 2% paraformaldehyde and 2% glutaraldehyde in PBS. Samples were embedded in the Embed-812 resin mixture and sectioned on an ultramicrotome (LKB Ultratome V; Leica, Paris, France) using a glass knife. Cross-sections of 0.27 μm thickness were stained with 1% methylene blue.

Axon cross section images were obtained using a Nikon Ti2 Eclipse microscope and NIS-Elements imaging software (Nikon). For each optic nerve cross section, 4 images obtained using a 60X (oil) objective were stitched in order to capture the entire nerve. This was performed on three cross sections per optic nerve, to ensure consistency. Axon count analysis was performed using the AxoNet plugin for ImageJ.^33^ The final axon count was averaged over the three cross-sections and then divided by the total optic nerve area to determine mean axon density per nerve.

The same mouse optic nerve specimens were thinly sectioned (60-80 nm) for transmission electron microscopy. Samples were collected on copper grids, counterstained with uranyl acetate and Sato’s lead, and then examined using an electron microscope (JEM-1400; JEOL) at 60 kV. Images were collected using a charge-coupled device camera (Orius; Gatan).

### Western Blot for GFAP Quantification

To quantify GFAP protein abundance, freshly dissected retinas were sonicated in lysis buffer [25 mM HEPES buffer, pH 7.4, 150 mM NaCl, 5 mM MgCl^2^, and protease inhibitors (Complete Mini, Roche, Indianapolis, IN) with 1% Triton X-100] and the protein concentration of each lysate was determined with a colorimetric assay (Bio-Rad). After mixing with SDS-PAGE sample buffer, four retinal lysates from each experimental group were separated on 10–20% SDS-PAGE gels, transferred onto polyvinylidene fluoride (PVDF) membranes and blotted with the indicated primary antibodies overnight. Membranes were washed in 0.05% Tween 20 and -actin, which served as the loading incubated with the appropriate secondary antibody for two hours. The Odyssey CLx imaging system (LiCor) was used to image and quantify band intensities. For each sample, the intensity of the GFAP band was normalized to that of the band for β control.

### Experimental Design and Statistical Analysis

Histological assessment of RGC soma and axon survival was performed on *Vglut2-Cre;ndufs4^loxP/loxP^* and littermate control mice with both sexes represented. In these experiments, 8-14 retinas or optic nerves were analyzed for each genotype and O_2_ concentration at each time point. Histological quantification of GFAP localization within Müller glia processes and Iba1-positive mononuclear cells in the inner retina was performed in 4 or 5 retinas for each genotype, O_2_ concentration, and time point. In all quantitative histological analyses, the observer was masked to the identity of each sample. For all experiments, statistical comparisons between groups were made with the Wilcoxon rank sum test to account for non-parametric data. All data analysis for this study was generated using SAS/STAT software, Version 9.4 of the SAS System for Windows (SAS Institute, Inc.). Data are presented graphically as mean ± SEM.

## RESULTS

### Hypoxia achieves complete histological rescue of RGCs in *Vglut2-Cre;ndufs4^loxP/loxP^* mice at P60 and remains therapeutic at P90

We have previously observed that RGC somas and axons develop normally in *Vglut2-Cre;ndufs4^loxP/loxP^* mice, with no histological phenotype at P30 and only mild degeneration observed at P45.^16^ Given the absence of early degeneration prior to weaning, we elected to initiate continuous hypoxia at P25. *Vglut2-Cre;ndufs4^loxP/loxP^* mice and control *ndufs4^loxP/loxP^* littermates lacking Cre recombinase were subjected to normoxia (21% O_2_) or hypoxia (11% O_2_) through P60. At this age, the conditional knockout mice did not manifest an overt neurological phenotype under either oxygen concentration, consistent with prior observations.^16,34^ RGC soma density was assessed on retinal flat mounts stained with the RGC marker RBPMS1 (Fig. 1A). As we had observed previously, *Vglut2-Cre;ndufs4^loxP/loxP^* mice raised entirely under normoxia demonstrated a reduction of RGC density by approximately one-third, and this was observed at locations proximal, intermediate, and distal from the optic nerve head (p≤0.01 for all locations; Fig. 1B). In contrast, no degeneration of RGC somas was observed when mice were treated with hypoxia; the soma density was indistinguishable between *Vglut2-Cre;ndufs4^loxP/loxP^* housed at 11% O_2_ from P25 to P60 and control *ndufs4^loxP/loxP^* mice exposed to either O_2_ concentration.

**Figure 1.**
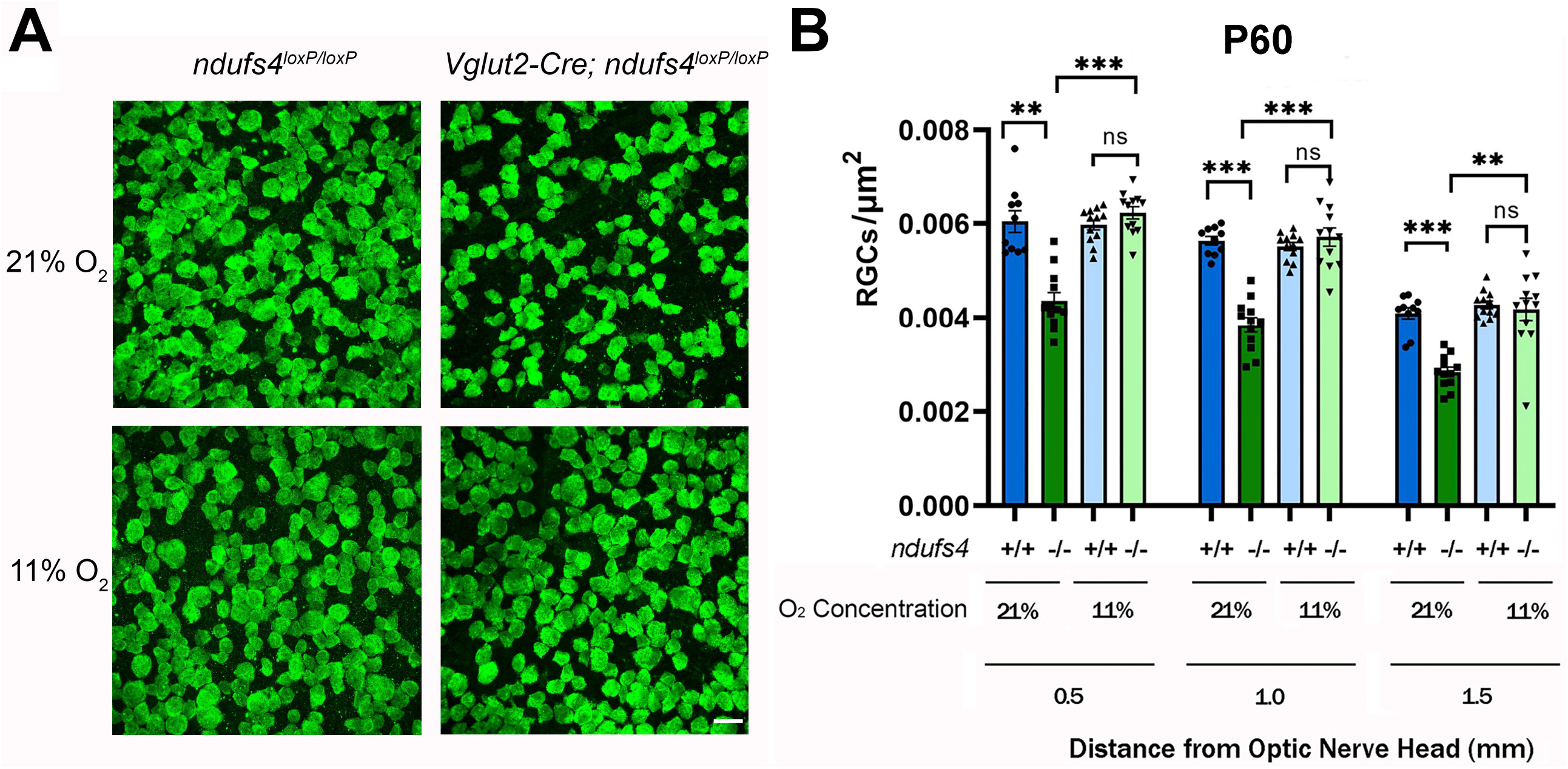
Continuous exposure to 11% O_2_ prevents RGC soma degeneration at P60 in *Vglut2-Cre;ndufs4^loxP/loxP^* mice. (A) Representative images obtained from retinal flat mounts at locations 1.0 mm from the optic nerve head, with RGC somas immunolabeled with RNA-Binding Protein 1 (RBPMS1; green). Control *ndufs4^loxP/loxP^* mice lacking Cre recombinase (left panels) and conditional knockout *Vglut2-Cre;ndufs4^loxP/loxP^* mice (right panels) were housed under normoxic conditions (21% O_2_; top row) or under hypoxia (11% O_2_; bottom row) from P25 to P60. The reduction of RGC soma density in normoxic *Vglut2-Cre;ndufs4^loxP/loxP^* retinas at P60 was prevented by continuous hypoxia. Bar, 20 μm. (B) RGC soma density (RGC somas/μm^2^) for the indicated genotypes and environmental O_2_ concentrations are shown at three different distances from the optic nerve head: 0.5, 1.0, and 1.5 mm. *ndufs4*^+/+^ indicates intact copies of both alleles (*ndufs4^loxP/loxP^* mice), whereas *ndufs4^-/-^*indicates deletion of both alleles (*Vglut2-Cre;ndufs4^loxP/loxP^* mice). Individual data points are depicted as black circles, squares, or triangles. Data are presented as mean ± SEM. Statistical comparisons between groups are indicated above the bars, with the following significance designations: ns, non-significant; **, p≤ 0.01; ***, p≤ 0.001.

To determine the durability of the hypoxia-mediated RGC rescue, additional cohorts of mice were analyzed at P90, the latest time point at which the normoxic *Vglut2-Cre;ndufs4^loxP/loxP^* mice could be assessed prior to death. At this time point, the conditional knockout mice maintained under normoxia were ataxic and manifested stiffness of the limbs, whereas those housed at 11% O_2_ had a grossly normal systemic phenotype. Further RGC loss of approximately 45% was observed in *Vglut2-Cre;ndufs4^loxP/loxP^* mice raised entirely under normoxia (Fig. 2A,B). In contrast to the earlier time point of P60, the rescue of RGC degeneration by hypoxia was no longer complete at P90; however, the extent of RGC soma loss was reduced by >50% at all three locations within the retina (p<0.001 for all). In this experiment, cohorts of heterozygous *Vglut2-Cre;ndufs4^loxP/+^* mice were also included, in order to verify that loss of only one copy of *ndufs4* in RGCs is aphenotypic and that expression of Cre recombinase itself had no effect on our analyses. Consistent with prior reports of a normal systemic phenotype in heterozygous germline *ndufs4^+/-^*mice,^30^ we observed that RGC soma and axon density were completely normal at P90 regardless of the ambient O_2_ concentration, confirming that this genotype may serve as a useful control in future experiments, as it allows for more efficient breeding of experimental mice.

**Figure 2.**
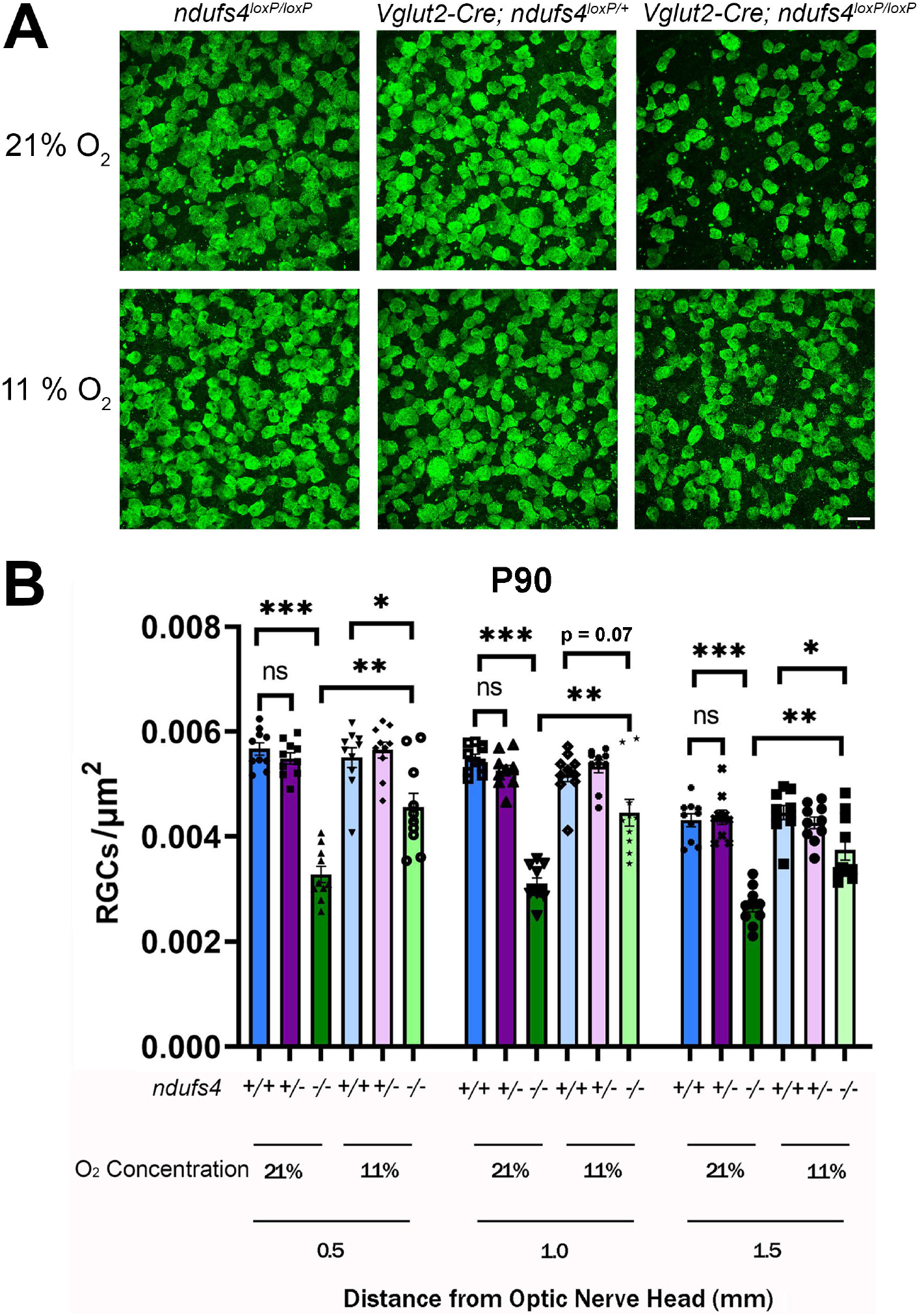
RGC soma degeneration in *Vglut2-Cre;ndufs4^loxP/loxP^* mice is partially rescued by hypoxia at P90. (A) Representative images obtained from retinal flat mounts at locations 1.0 mm from the optic nerve head, with RGC somas immunolabeled with RBPMS1 (green). Control *ndufs4^loxP/loxP^* mice lacking Cre recombinase (left panels), *Vglut2-Cre;ndufs4^loxP/+^* mice with only one allele of *ndufs4* deleted by Cre (middle panels), and *Vglut2-Cre;ndufs4^loxP/loxP^* mice with both alleles of *ndufs4* deleted (right panels) were housed under normoxic conditions (21% O_2_; top row) or under hypoxia (11% O_2_; bottom row) from P25 to P90. Only with deletion of both copies of *ndufs4* is RGC soma degeneration observed at P90, and this level of degeneration is markedly reduced by continuous hypoxia. Bar, 20 μm. (B) RGC soma density (RGC somas/μm^2^) for the indicated genotypes and environmental O_2_ concentrations are shown at three different distances from the optic nerve head: 0.5, 1.0, and 1.5 mm. *ndufs4*^+/+^ indicates intact copies of both alleles (*ndufs4^loxP/loxP^* mice; blue bars), *ndufs4^+/-^*indicates deletion of one allele of *ndufs4* (*Vglut2-Cre;ndufs4^loxP/+^* mice; purple bars), and *ndufs4^-/-^*indicates deletion of both alleles (*Vglut2-Cre;ndufs4^loxP/loxP^* mice; green bars). Individual data points are overlayed on each bar. Statistical comparisons between groups are indicated above the bars, with the following significance designations: ns, non-significant; *, p≤0.05; **, p≤0.01; ***, p≤ 0.001.

### Continuous hypoxia reduces axonal degeneration in *Vglut2-Cre;ndufs4^loxP/loxP^* mice

As a complementary assessment of the neuroprotective effect of hypoxia on RGCs from *Vglut2-Cre;ndufs4^loxP/loxP^* mice, optic nerve cross-sections were obtained in order to quantify RGC axon density and to assess for morphological rescue. At P60 axon density was reduced by 58% in the optic nerves of normoxic *Vglut2-Cre;ndufs4^loxP/loxP^* mice compared to *Vglut2-Cre;ndufs4^loxP/+^* control mice, whereas the axon density was significantly increased in the knockouts raised under hypoxia (p<0.001) and no different from the controls (Fig. 3A,B). Electron microscopy revealed reduced axon density with increased surrounding fibrosis and abnormal myelination patterns in the optic nerves of normoxic *Vglut2-Cre;ndufs4^loxP/loxP^* mice (Fig. 3C). In contrast, an orderly, healthy appearance of myelinated RGC axons was observed in hypoxic *Vglut2-Cre;ndufs4^loxP/loxP^* mice, indistinguishable from that of control littermates.

**Figure 3.**
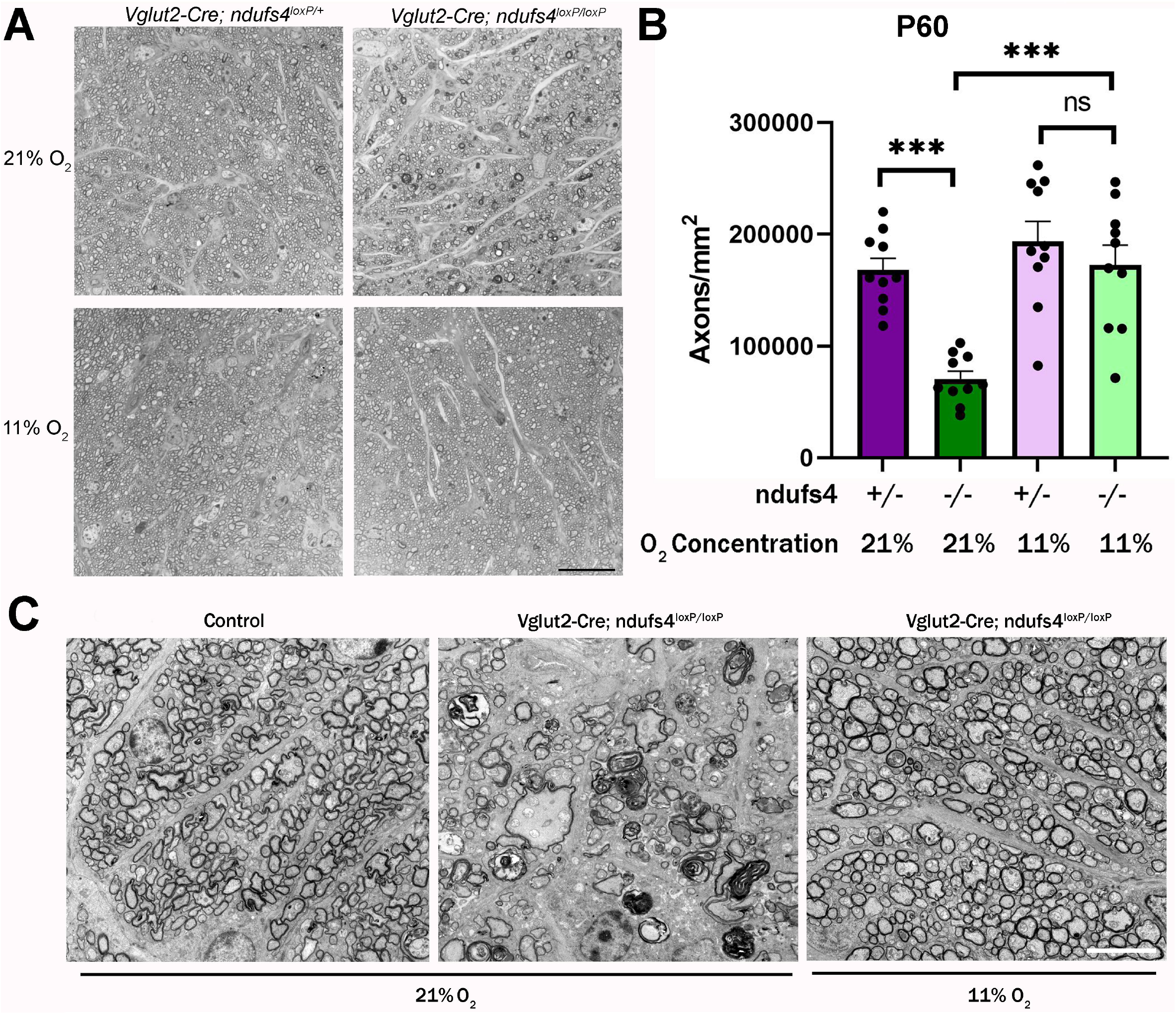
RGC axonal degeneration in *Vglut2-Cre;ndufs4^loxP/loxP^* mice at P60 is prevented by continuous hypoxia. (A) Representative light micrographs of optic nerve cross sections stained with methylene blue. Control *Vglut2-Cre;ndufs4^loxP/+^* mice (left column) and conditional knockout *Vglut2-Cre;ndufs4^loxP/loxP^* mice (right column) were housed under normoxia (21% O_2_, upper row) or hypoxia (11% O_2_, lower row) from P25 to P60. Bar, 20 μm. (B) RGC axon densities in optic nerve cross sections determined using AxoNet automated axon quantification. *ndufs4^+/-^*indicates deletion of one allele of *ndufs4* (*Vglut2-Cre;ndufs4^loxP/+^* mice; purple bars), and *ndufs4^-/-^*indicates deletion of both alleles (*Vglut2-Cre;ndufs4^loxP/loxP^* mice; green bars). The O_2_ concentration is indicated for each group. Individual data points are depicted as black circles. Data are presented as mean ± SEM. Statistical comparisons between groups are indicated above the bars, with the following significance designations: ns, non-significant; ***, p≤0.001. (C) Electron micrographs (5000X) of optic nerve cross sections at P60 demonstrate preservation of normal axon morphology and abundance in *Vglut2-Cre;ndufs4^loxP/loxP^* mice raised under continuous hypoxia (right) compared to normoxia (middle). Bar, 5 μm.

Consistent with our analysis of RGC somas, a partial rescue of RGC axons was apparent at P90. While the hypoxia-treated *Vglut2-Cre;ndufs4^loxP/loxP^* mice exhibited a 32% reduction in axon density compared to controls mice, this represented a rescue of 45% of the axons lost under normoxic conditions (Fig. 4A,B). Ultrastructural analysis of the optic nerves via electron microscopy demonstrated the interim development of abnormal myelination patterns in the hypoxia-raised *Vglut2-Cre;ndufs4^loxP/loxP^* mice at P90 compared to the P60 time point; however, the morphological abnormalities were demonstrably less severe than in the cohort kept continuously under normoxia (Fig. 4C).

**Figure 4.**
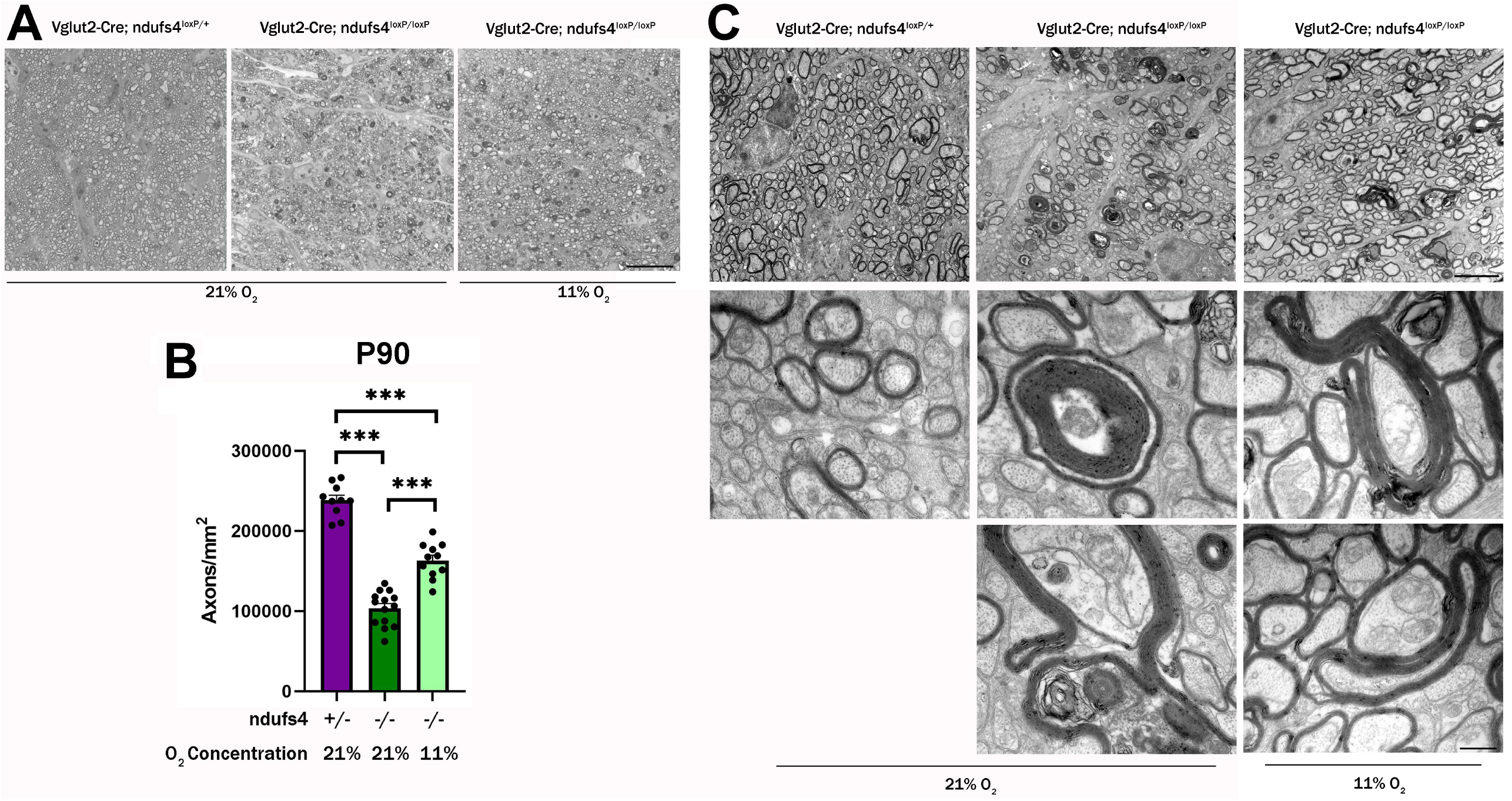
Retinal ganglion cell axonal degeneration is reduced in P90 *Vglut2-Cre;ndufs4^loxP/loxP^* mice treated with hypoxia. (A) Representative light micrographs of optic nerve cross sections stained with methylene blue for *Vglut2-Cre;ndufs4^loxP/+^* control mice kept under normoxia (21% O_2_, left), *Vglut2-Cre;ndufs4^loxP/loxP^* mice kept under normoxia (middle), and *Vglut2-Cre;ndufs4^loxP/loxP^* mice treated with hypoxia (11% O_2_, right) from P25 to P90. Bar, 20 μm. (B) RGC axon densities in optic nerve cross sections at P90 determined using AxoNet automated axon quantification. *ndufs4^+/-^*indicates deletion of one allele of *ndufs4* (*Vglut2-Cre;ndufs4^loxP/+^* mice; purple bar), and *ndufs4^-/-^*indicates deletion of both alleles (*Vglut2-Cre;ndufs4^loxP/loxP^* mice; green bars). The O_2_ concentration is indicated for each group. Individual data points are depicted as black circles. Data are presented as mean ± SEM. Statistical comparisons between groups are indicated above the bars: ***, p≤ 0.001. (C) Electron micrographs with magnifications of 5,000X (top row; bar, 5 μm) and 40,000X (bottom two rows; bar 0.5 μm) demonstrate axon density and morphology in P90 optic nerve cross sections from normoxic *Vglut2-Cre;ndufs4^loxP/+^* control mice (left column) and *Vglut2-Cre;ndufs4^loxP/loxP^* mice kept under normoxia (middle column) or hypoxia (right column). The higher magnification images demonstrate typical abnormalities in myelination that occur in *Vglut2-Cre;ndufs4^loxP/loxP^* optic nerves, including thickening and doubling of the myelin sheaths and incomplete enclosure of axons. While these abnormalities may be observed in hypoxia-treated animals (right), they are less frequent and generally less severe.

### Hypoxia partially reduces retinal neuroinflammation in *Vglut2-Cre;ndufs4^loxP/loxP^* retinas

A recent study in germline *ndufs4^-/-^*mice found evidence that reducing neuroinflammation by depleting leukocytes had a dramatic therapeutic effect, prolonging lifespan and reducing neurologic dysfunction in the mice.^35^ Notably, it has also been shown in the germline *ndufs4^-/-^*mouse^36^ and in our *Vglut2-Cre;ndufs4^loxP/loxP^* conditional knockout mouse line^16^ that reactive gliosis and inner retinal accumulation of Iba1-positive mononuclear inflammatory cells occur alongside RGC degeneration. Given the neuroprotective effect of hypoxia on *ndufs4*-deficient RGCs, we wondered whether continuous hypoxia would produce a similar reduction of retinal neuroinflammation in *Vglut2-Cre;ndufs4^loxP/loxP^* mice. As before, the *Vglut2-Cre;ndufs4^loxP/loxP^* mice and control *Vglut2-Cre;ndufs4^loxP/+^* littermates were exposed to either 21% O_2_ or 11% O_2_ from P25 until harvesting of ocular tissue at P60 or P90. Reactive gliosis was determined by assessing for up-regulation of glial fibrillary acidic protein (GFAP), an intermediate filament expressed constitutively by retinal astrocytes and detectable in Müller glia as a response to retinal pathology. At P60, the abundance of GFAP protein was increased in the retinas of *Vglut2-Cre;ndufs4^loxP/loxP^* mice raised under normoxia by four-fold compared to *Vglut2-Cre;ndufs4^loxP/+^* controls (p<0.05), whereas the retinal GFAP protein level in hypoxic *Vglut2-Cre;ndufs4^loxP/loxP^* mice was not increased (Fig. 5A). Consistently, while radial localization of up-regulated GFAP in Müller cells was not observed in control *Vglut2-Cre;ndufs4^loxP/+^* mice, it was readily identified in retinal sections obtained from *Vglut2-Cre;ndufs4^loxP/loxP^* mice raised under normoxia, but not hypoxia at P60 (Fig. 5B). Similar to the partial hypoxia-mediated rescue of RGC somas seen at P90, GFAP up-regulation was not completely prevented at this time point but was reduced by two-fold (p<0.05) (Fig. 5C). The number of GFAP-positive Müller radial processes was also intermediate in P90 *Vglut2-Cre;ndufs4^loxP/loxP^* mice treated with hypoxia compared to controls and *Vglut2-Cre;ndufs4^loxP/loxP^* mice raised under normoxia (Fig. 5D).

**Figure 5.**
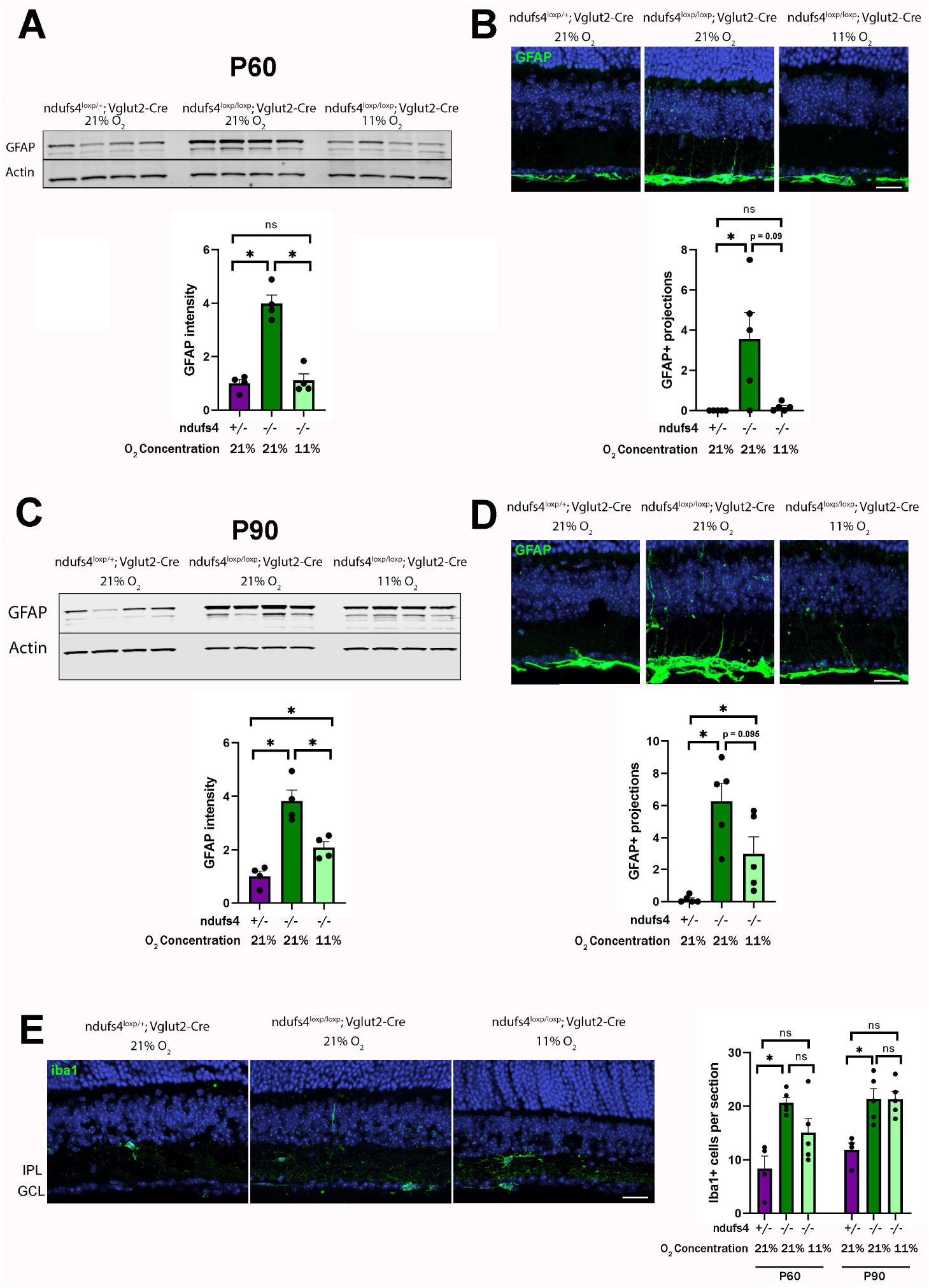
The effect of continuous hypoxia on retinal neuroinflammation in *Vglut2-Cre;ndufs4^loxP/loxP^* mice. (A) Glial fibrillary acidic protein (GFAP) expression levels in P60 retinal lysates assessed by Western blot, with four replicates each for normoxic control *Vglut2-Cre;ndufs4^loxP/+^* mice (left) and for *Vglut2-Cre;ndufs4^loxP/loxP^* mice exposed to normoxia (middle) or hypoxia (right). The graph below depicts the relative increase in GFAP band intensity normalized to the actin compared to the control group. (B) Representative images of P60 retinal cross sections immunolabeled with GFAP (green). In control *Vglut2-Cre;ndufs4^loxP/+^* mice (left), GFAP signal is present only in astrocytes of the inner-most retina, whereas in normoxic *Vglut2-Cre;ndufs4^loxP/loxP^* mice (middle), GFAP-positive radial projections of Müller glia are observed. This reactive gliosis is prevented by treating the mice with hypoxia (right). The graph below depicts the mean number of GFAP-positive Muller projections identified per 100-μm segment of retina for each group. (C) Western blot assessment of GFAP protein levels in retinal lysates at P90. As in Panel A, the normalized GFAP band intensity is plotted relative to control. (D) Representative images of retinal cross sections immunolabeled with GFAP (green) at P90 for the same experimental groups depicted in Panel B. The abundance of GFAP-positive radial Müller projections is depicted in the graph below. (E) P60 retinal cross sections were labeled with Iba1 (green) to identify mononuclear inflammatory cells. Bar, 20 μm. The mean number of Iba1-positive cells within the ganglion cell layer (GCL) or inner plexiform layer (IPL) across a retinal section are depicted in the graph to the right for retinas obtained at P60 and P90. For all graphs, bars depict mean ± SEM, with individual data points displayed; ns, not significant; *, p<0.05.

The effect of hypoxia on inner retinal Iba1-positive mononuclear cell accumulation in *Vglut2-Cre;ndufs4^loxP/loxP^* mice was less pronounced. As we previously observed,^16^ there was a significant >2-fold increase in Iba1-positive cell abundance in P60 *Vglut2-Cre;ndufs4^loxP/loxP^*retinas under normoxia compared to controls; however, an intermediate abundance was observed at this early time point in mutant mice raised under hypoxia that was not significantly lower than those raised under normoxia (Fig. 5E). By P90, mononuclear cell accumulation in the inner retina in *Vglut2-Cre;ndufs4^loxP/loxP^* mice was identical regardless of O_2_ concentration. Our observations suggest that RGC dysfunction in the setting of complex I deficiency may induce an early inflammatory response that is anticipatory to overt RGC death.

## DISCUSSION

In this study, we have shown that continuous hypoxia is neuroprotective when applied prior to the onset of RGC degeneration in mice with RGC-specific deletion of *ndufs4*. With hypoxia treatment, no discernible loss of RGC somas or axons was observed at P60, a time point at which approximately one-third of RGCs have degenerated in *Vglut2-Cre;ndufs4^loxP/loxP^* mice raised entirely under normoxia. Compared to the systemic phenotypes of germline *ndufs4^-/-^*mice (e.g. ataxia, stiffness, inactivity, weight loss, early death), which have been reported to be delayed by a half-year or more by hypoxia,^28,29^ the therapeutic effect on optic atrophy in *Vglut2-Cre;ndufs4^loxP/loxP^* mice is more time-limited, as the 100% rescue of RGCs observed at P60 diminished to approximately 50% after one additional month, with accompanying signs of reactive gliosis by astrocytes and Müller cells.

Another interesting finding in our study is a trend toward an increase in mononuclear inflammatory cell accumulation in the inner retinas of *Vglut2-Cre;ndufs4^loxP/loxP^* mice at P60 despite 100% RGC rescue by hypoxia. As the complex I deficiency in this mouse model is intrinsic to RGCs, it seems likely that dysfunctional RGCs release proinflammatory signals prior to their degeneration. Indeed, a previous global gene expression analysis of the retinas of germline *ndufs4^-/-^*mice at P33 (prior to the onset of RGC degeneration) found inflammatory and immune-related pathways to be the most highly up-regulated compared to control retinas.^36^ Extrapolating from the significant benefits of inflammatory cell depletion on the neurological phenotype of germline *ndufs4^-/-^*mice,^35^ it may be that the retinal accumulation of mononuclear inflammatory cells (which has become maximal by P90 even with hypoxia treatment) exacerbates the degeneration of complex I-deficient RGCs. This raises the possibility that hypoxia and anti-inflammatory interventions might have additive or even synergistic therapeutic effects. Alternatively, it could be the case that the neuroprotective effect of hypoxia that we observed is entirely mediated by a reduction in neuroinflammation.

Regarding the more moderate reduction of optic atrophy by hypoxia in *Vglut2-Cre;ndufs4^loxP/loxP^* mice compared to the much more prolonged systemic improvement in germline *ndufs4^-/-^*mice, the most likely explanation is an exquisite sensitivity of RGCs to chronic complex I dysfunction. This notion is supported by the well-documented observation that, in the setting of the milder insults to complex I function associated with LHON, most patients develop optic atrophy in the absence of other neurological or systemic symptoms.^37^ In this context, the time-limited therapeutic effect of hypoxia on *ndufs4*-deficient RGCs may still hold significant relevance to developing treatments for LHON, as hypoxia-mediated neuroprotection of RGCs with milder complex I dysfunction could prove more durable. An important next step, therefore, would be to conduct a long-term assessment of the effect of hypoxia on the more slowly-developing optic neuropathies characteristic of the mouse models with LHON-associated mutations in complex I subunits.^12,13^

A longer-term goal will be to elucidate the mechanism(s) underlying the salutary effect of hypoxia on complex I-deficient RGCs. This might involve a decrease in cellular oxidative stress via a generalized reduction of the availability of molecular oxygen at the tissue level.^30,38^ Alternatively, RGC neuroprotection by hypoxia may arise from modulation of cellular metabolism or other features of mitochondrial biology due to activation of specific molecular pathways as an adaptation to hypoxia. A mechanistic understanding of this process could ultimately lead to targeted pharmacological interventions that could protect RGCs with mitochondrial impairment while potentially obviating the need for direct hypoxia.

In summary, we have demonstrated robust *in vivo* neuroprotection of complex I-deficient RGCs by continuous exposure to hypoxia. Our observations indicate that identifying the relevant cellular processes modulated by hypoxia may represent a critical step in addressing the unmet need of developing effective therapies for mitochondrial optic neuropathies like LHON.

## ACKNOWLEDGEMENTS

The authors would like to thank Vadim Arshavsky for critical review of the manuscript.

